# Melanin found in wheat spike husks

**DOI:** 10.1101/2023.05.30.542842

**Authors:** Mikhail S. Bazhenov, Dmitry Y. Litvinov, Mikhail G. Divashuk

## Abstract

Melanin is the dark polymer pigment found in all kingdoms of life. Plant melanin, formed through the oxidation and polymerization of phenolic compounds, does not contain nitrogen, however it possesses similar properties with melanin of animal, fungal or bacterial origin. Melanin in plants is usually found in seed coats or fruit pericarp and is ascribed mechanical barrier or some other protective functions. Wild and formerly locally cultivated wheat species, like Persian wheat (*Triticum carthlicum* Nevski) frequently exhibit black color of spike husks and awns. The pigment causing it and the biological purpose of this coloration was not clarified before. In this paper using standard chemical extraction procedures for anthocyanins and melanin, ultraviolet-visible-near-infrared spectroscopy and hyperspectral imaging, we prove that the black color of Persian wheat spikes is caused by melanin but not anthocyanins. Also, we show that the dark pigment in husks and awns is located in epiderma and subepidermal sclerenchyma cells, that implies melanin potentially to have mechanical-enhancing and protection function. Other possible functions of melanin in cereals are discussed.

## Introduction

Melanin is the black or dark brown polymer pigment found in all kingdoms of life, that does not have definite chemical structure. The plant melanin, also classified as allomelanin, is formed through the oxidation and polymerization of phenolic compounds. Catechol, caffeic, chlorogenic, protocatechuic, and gallic acids are considered to be the possible precursors of melanin in plants, while animal, fungi and bacterial melanin pigments (eumelanin and pheomelanin) are derivatives of tyrosine. Unlike animal, fungi or bacterial pigments, the plant melanin, or otherwise called phytomelanin, does not contain nitrogen, however resembles physiochemical properties of other melanin pigments [1].

The role of melanin in plants is not yet completely understood. Most probably it plays a role of a barrier-making substance in seed coats or fruit pericarp, that protects seeds from being eaten by insects or infected by fungi [2]. Formerly, phytomelanin was mainly known in Asteraceae and Asparagaceae families as a component of pericarp or seed coat of these plants [3]. For example, most agronomists are well familiar with phytomelanin layer in sunflower (*Helianthus annuus* L.) achenes pericarp, that confers resistant to larvae of the sunflower moth (*Homoeosoma electellum* Hulst) [4]. Now phytomelanin was found to be widespread in plants. For example, well-known enzymatic browning reaction that occurs in wounded plant tissues leads to formation of melanin. This reaction occurs when, due to the disruption of cells, the chloroplast-located polyphenol oxidases (PPOs) are released and interact with vacuolar substrates to produce o-quinones, which then polymerize to melanin. Melanin formation in intact plant tissues also probably involves PPOs [5], however other enzymes, like cell wall-associated laccases, are supposed to participate in extracellular melanin deposition [1]. Melanin was found in seed coats and fruit pericarps of a wide variety of plant families and species: watermelon, buckwheat, grape, tomato, fragrant olive, night jasmine, sesame, ipomoea, black mustard and rape, chestnut, garlic [1]. Recently, phytomelanin was found in cereals – in barley varieties possessing black lemmas or pericarp [6] and in black oat [7]. In sunflower and *Echinacea* achenes the melanin accumulates extracellularly between the hypodermis and sclerenchyma in the pericarp [2,4], in barley it forms small granules within chloroplasts of pericarp, lemma and palea [8], while in persimmon skin it is deposited on the cell walls of the upper epidermis and subepidermal cells [5]. Dark-colored barley seeds have higher contents of phenolic compounds and lignin than uncolored ones. It was suggested that melanin biosynthesis genes may be connected to phenylpropanoid-derived biosynthesis pathways [9].

Melanin from various sources finds numerous potential applications in cosmetics, agriculture, medicine and technology [10–12]. Melanin possesses properties of free radicals’ scavenger, metals chelating agent, ultraviolet- and radioactive-protecting properties [13–15]. In medicine it could be used as anti-inflammatory, immunostimulatory, digestive system protective and liver protective and antivenin remedy [16–22]. Thus, new sources of melanin can be of interest.

Some wild and formerly locally cultivated wheat species, like Persian wheat (*Triticum carthlicum* Nevski) exhibit black color of spike husks and awns. The biological purpose of this coloration is not understood, so it is not clear whether there are any potential benefits from introducing this trait into wheat cultivars. Since the nature of the black color of wheat spikes remains unclear [23], we decided to clarify the type of the pigment that provides this color. In this study, we tested the hypothesis of phytomelanin to be the cause of the black color of Persian wheat spikes.

## Materials and methods

### Plant material

In order to test the hypotheses on the presence of melanin in black-colored wheat husks, we took an accession of Persian wheat Line 5999 (*Triticum carthlicum* Nevski, VIR^1^ catalog #26828, collected in 1930 near village Kumukh, Dagestan, Russia). The wheat plants were grown in Moscow Oblast (WGS 84 coordinates 55.82106, 38.22563) in 2022. The mature spikes (**Figure 1a**) were threshed manually, and the husks were gathered. Sunflower (*Helianthus annuus* L.) black-colored achenes husks were taken as a positive melanin-containing control [24], while dark-colored blueberry (*Vaccinium corymbosum* L.) fruit exocarp, red-colored and black-colored tissues of tulip (*Tulipa* L., Darwin hybrid ‘Apeldoorn’) perianth [25] were taken as anthocyanin-positive controls (**Figure 1**). The tulip was grown at the same place as wheat, while sunflower and blueberry fruits were purchased in local supermarket.

**Figure 1.**
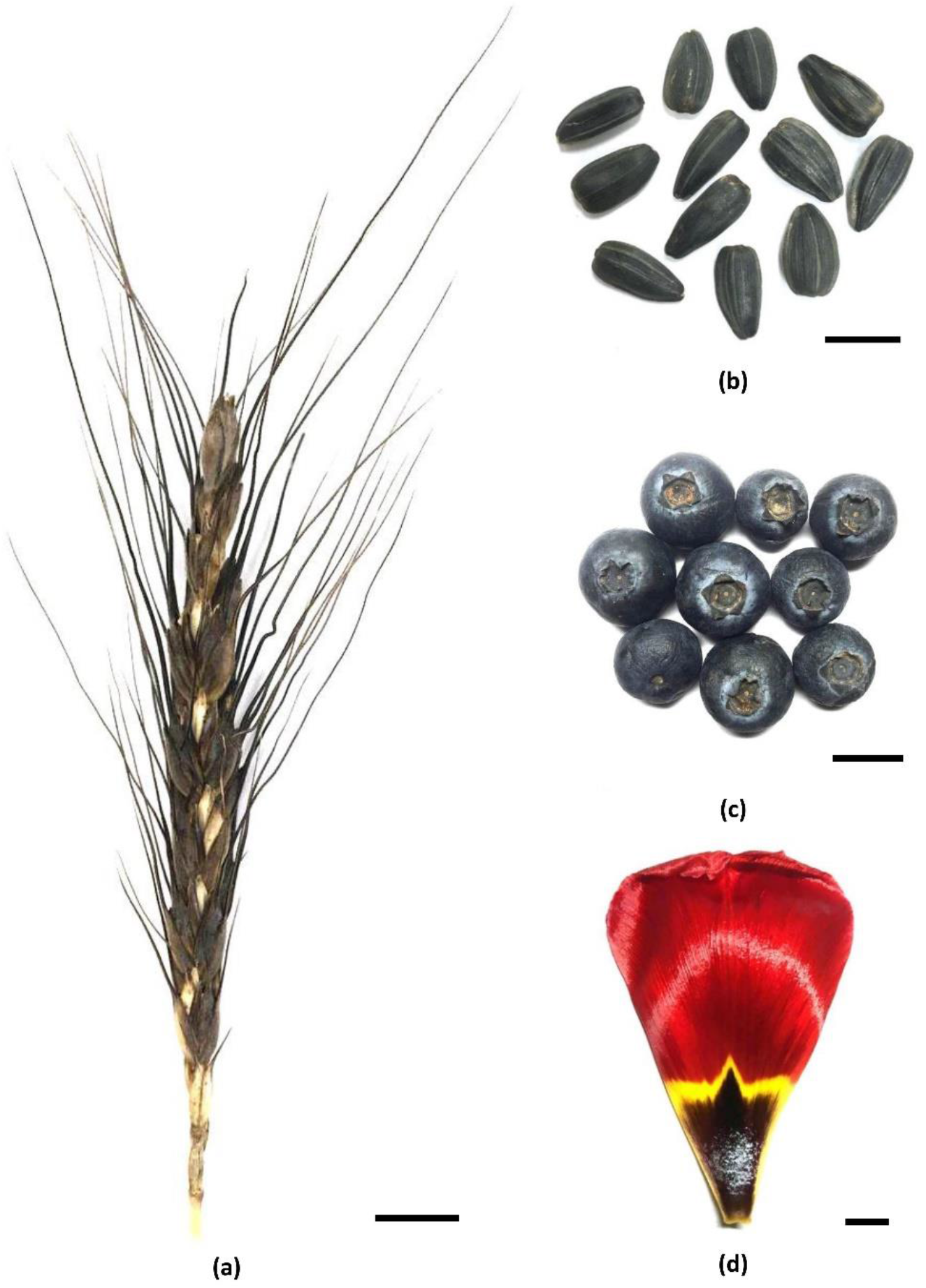
The plant material used for pigment extraction: the Persian wheat Line 5999 spike **(a)**, sunflower achenes **(b)**, blueberry fruits **(c)**, tulip perianth segments **(d)**. The horizontal bar in the lower right corner of each image is 1 cm long.

### Preparation of melanin and anthocyanin extracts

About 100 mg of plant tissues, fresh (tulip perianth, blueberry exocarp) or dry (wheat and sunflower husks) were placed in 2 ml Eppendorf centrifuge tubes. The extraction solutions described below were added to fresh tissues before grinding. The grinding was done using 7 mm stainless steel beads with TissueLyser II bead mill (Qiagen, Hilden, Germany), 2 minutes totally at 25 Hz. The extraction solutions were added before the grinding in case of fresh samples, or after the grinding in case of dry samples.

To extract melanin, 1 ml of 0.3 M NaOH aqueous solution [24] was added to grinded tissues per tube, while for extraction of anthocyanins, 95% ethanol with addition of 0.1 M citric acid [26] was used in the same quantity. The contents of the tubes were shaken on a TissueLyser II 30 seconds at 25 Hz and then incubated on a tube rotator for 30 minutes at room temperature. After incubation, the tubes were centrifuged at 5000 g, and the supernatant was transferred to fresh tubes. The photograph of the tubes was taken using a smartphone camera.

### Measurement of UV-Visible absorption spectra

The absorbance spectrum of extracts was measured on a SPECTROstar Nano (BMG LABTECH, Germany) spectrometer in the UV, visible and near infrared regions (220-1000 nm) with 1 nm resolution. The measurements were done in 1 cm quartz cuvette against the corresponding extraction buffer as a blank. After the first measurement, the extracts were diluted to adjust the absorbance to about 1 OD at 400 nm. The spectral data were exported to Excel, recalculated to obtain 1 OD at 400 nm and plotted as *log*_*10*_ values of the absorbance to compensate strong differences in spectra between the samples.

### Hyperspectral imaging and reflection spectra in visible and near infrared regions

Hyperspectral imaging in visible and near infrared regions (400-1000nm) with 5 nm resolution was done using Muses9-HS hyperspectral camera (Spectricon, Greece) with its accessory light source (LS) combining halogen lamp and LED for UV and blue spectral regions. Spectralon diffuse reflectance standard (Labsphere – Halma,PLC, UK) was placed near the samples and used to calculate the reflectance of the samples as the ratio of the intensity of a sample and the intensity of the Spectralon.

### Histology of wheat spike

The lemma scale of Persian wheat spire was cast in paraffin block and cross-section cuts were made using hand microtome and razor blade. The cross-sections were placed in 70% ethanol solution on microscope slides and visualized using the Biolam D-11 microscope (LOMO, St. Petersburg, Russia) equipped with color digital camera. No chemical staining was applied.

### Data and image processing

The spectral plots were built in MS Excel 2016. Hyperspectral images were analyzed using Muses9HS software. The photo images were adjusted in Adobe Photoshop CS6.

## Results

### The reflectance spectrum of wheat lemma is similar to the spectrum of sunflower achene

The reflectance of dark-colored wheat spike lemma, sunflower achene, blueberry fruit and dark-colored part of tulip perianth showed very low values in the entire visible region of the spectrum and in the initial part of the near infrared region up to 725 nm, what was perceived by the eye as almost black. However, the specimens differed significantly in near infrared spectral region from 740 to 1000 nm (**Figure 2**). The reflectance of sunflower achenes was uniformly low in all range of wavelength from 400 to 1000 nm. Wheat light-colored lower part of lemma showed linear rise of reflectance in all the measured range. The dark-colored upper part of lemma showed slow gradual increase of reflectance only in infrared part of spectrum in the range from 700 to 1000 nm. The blueberry fruit and the dark part of tulip perianth, unlike dark wheat tissue, showed rapid raise of reflectance in the interval between 710 and 860 nm. Blueberry showed maximum reflectance at about 900 nm, while tulip dark part of perianth showed maximum at 1000 nm. The red part of tulip perianth showed low reflectance below 600 nm with a rapid rise above this wavelength.

**Figure 2.**
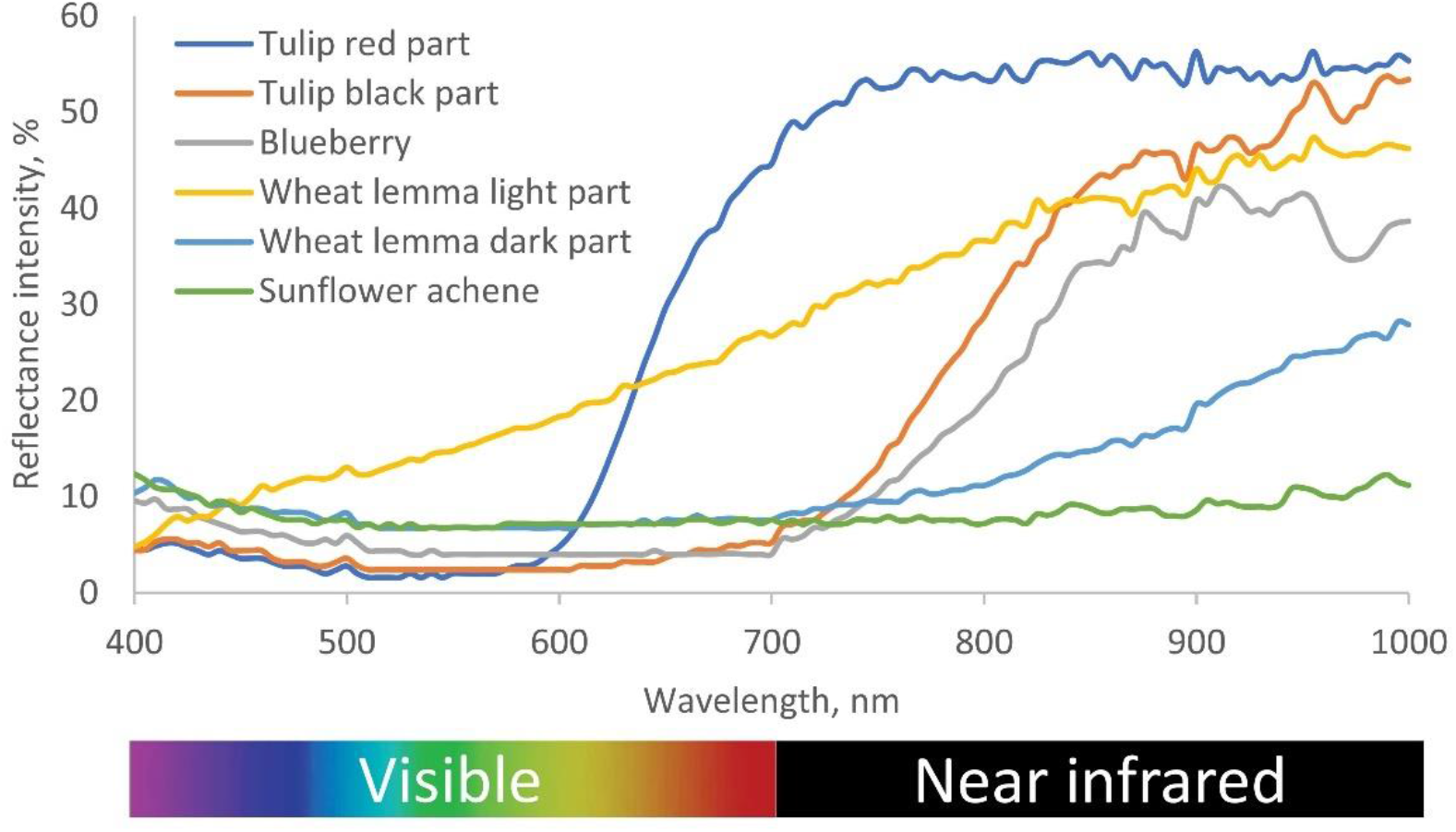
The reflectance spectra obtained from intact samples using hyperspectral camera.

### The solubility and pigment spectrum of dark colored Persian wheat husks suggest the presence of melanin rather than anthocyanins

The sodium hydroxide solution produced colored extracts for each plant tissue sample (**Figure 3**). The extract was yellow for the dark part of tulip perianth, brown for red part of one, and dark-brown for blueberry fruit skin, sunflower pericarp and wheat dark-colored husks. The acidified alcohol produced colored extracts only for anthocyanin-containing tissues. It was magenta for the dark part of tulip perianth, scarlet for red part of tulip perianth, and eggplant color for blueberry skin. For sunflower pericarp the alcohol extract was colorless and almost colorless pale-yellow for wheat husks.

**Figure 3.**
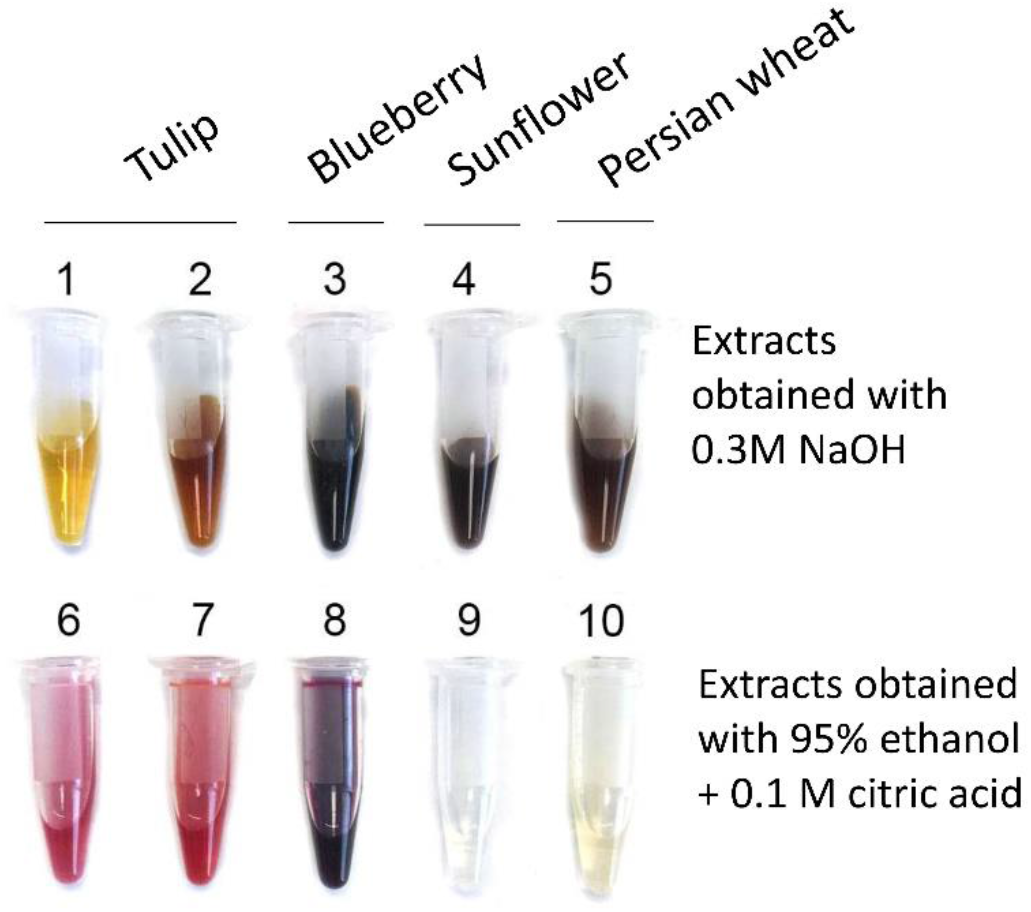
Plant tissue extract obtained with 0.3 M NaOH (1 – 5) or 95% ethanol with 0.1 M citric acid (6 – 10). Tulip black perianth tissue (1, 6), tulip red perianth tissue (2, 7), blueberry exocarp (3, 8), sunflower achenes pericarp (4, 9), Persian wheat dark-color spike husks (5, 10).

The spectrum measurements of the extracts showed that wheat and sunflower have much more similarity than wheat and tulip or wheat and blueberry for both sodium hydroxide (alkaline) and alcohol extracts. Both sunflower and wheat alkaline extracts showed the spectrum with maximum absorbance in ultraviolet, minimum in infrared and gradual exponential decrease between these marginal wavelengths point of the spectrum (**Figure 4a**). Tulip alkaline extracts were a bit different from sunflower and wheat, showing some peaks in ultraviolet (UV) and visible (VIS) light. Blueberry alkaline extract had some similarities to melanin-containing sunflower extract.

**Figure 4.**
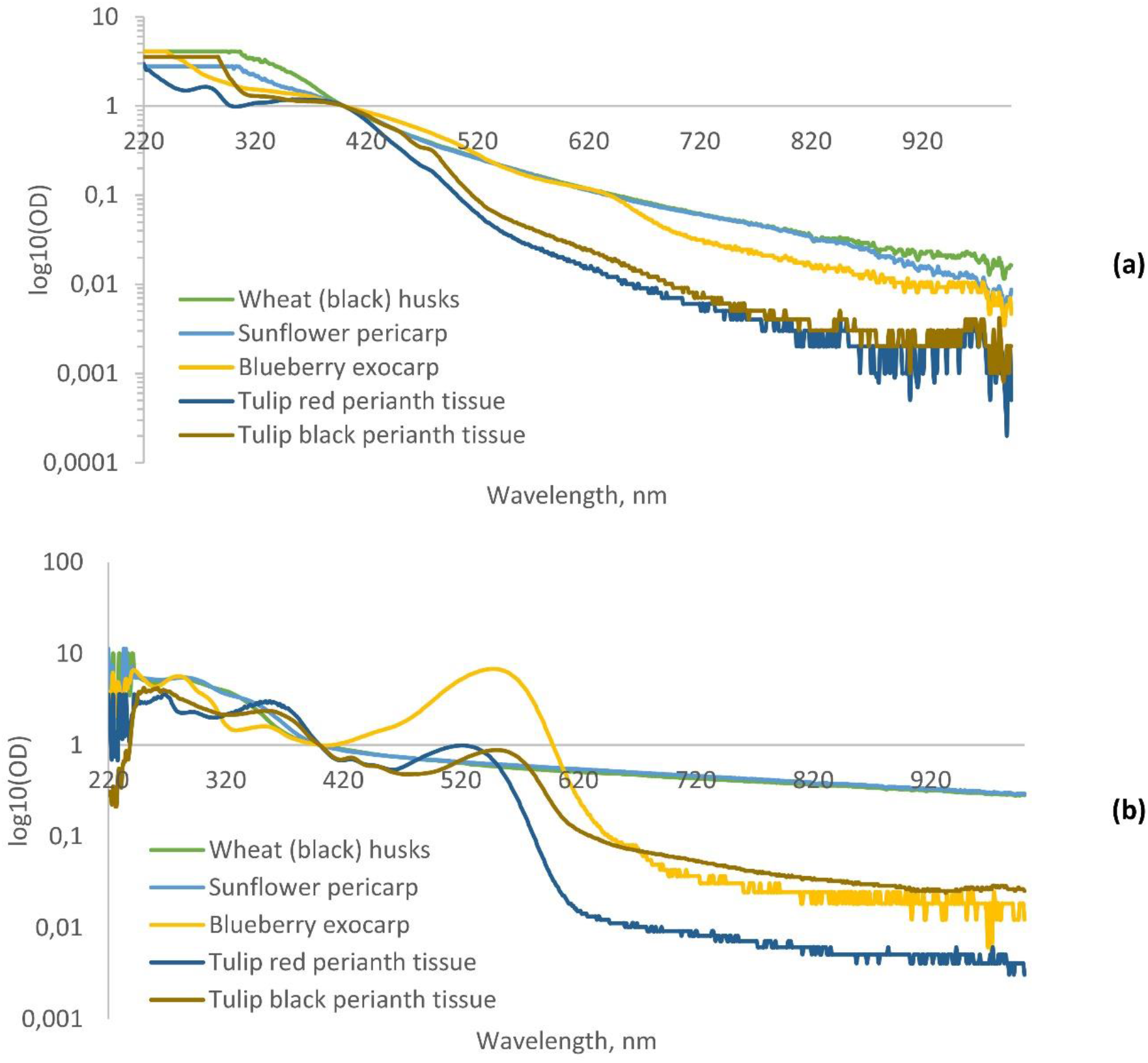
Spectra of absorbance of plant tissue extracts obtained with 0.3 M NaOH **(a)** or 95% ethanol with 0.1 M citric acid **(b)**, adjusted to get optical density (OD) equal to 1 at 400 nm, measured against corresponding blank solvent.

Alcohol extracts of blueberry and dark part of tulip perianth showed high peaks of absorbance in visible range of spectrum at about 547-550 nm, while red tulip perianth – at about 521 nm (**Figure 4b**). A small peak of absorbance at 427 nm was specific to tulip. In UV part of the spectrum, a peak at 354 nm was present for all anthocyanin-containing tissues. Blueberry also showed the peaks of absorbance at 281 and 242 nm, while tulip – in a region of 246-268 nm. For wheat and sunflower alcohol extracts, a single clear peak at 283-284 nm was observed in the UV part of spectrum, followed by gradual exponential decrease of absorbance towards infrared part.

### Micro-anatomical distribution of pigments in Persian wheat dark-color spike husks

Further we conducted anatomy study to reveal pigment localization in Persian wheat spike tissues (**Figure 5**). On the cross-sections of lemma of mature spike and its awn it is visible, that dark pigment is accumulated mainly inside epidermal cells, including sometimes trichomes, and in underlying 1-2 layers of sclerenchyma cells. The pigment is most concentrated in the center of the cells and is also present in cell walls of sclerenchyma.

**Figure 5.**
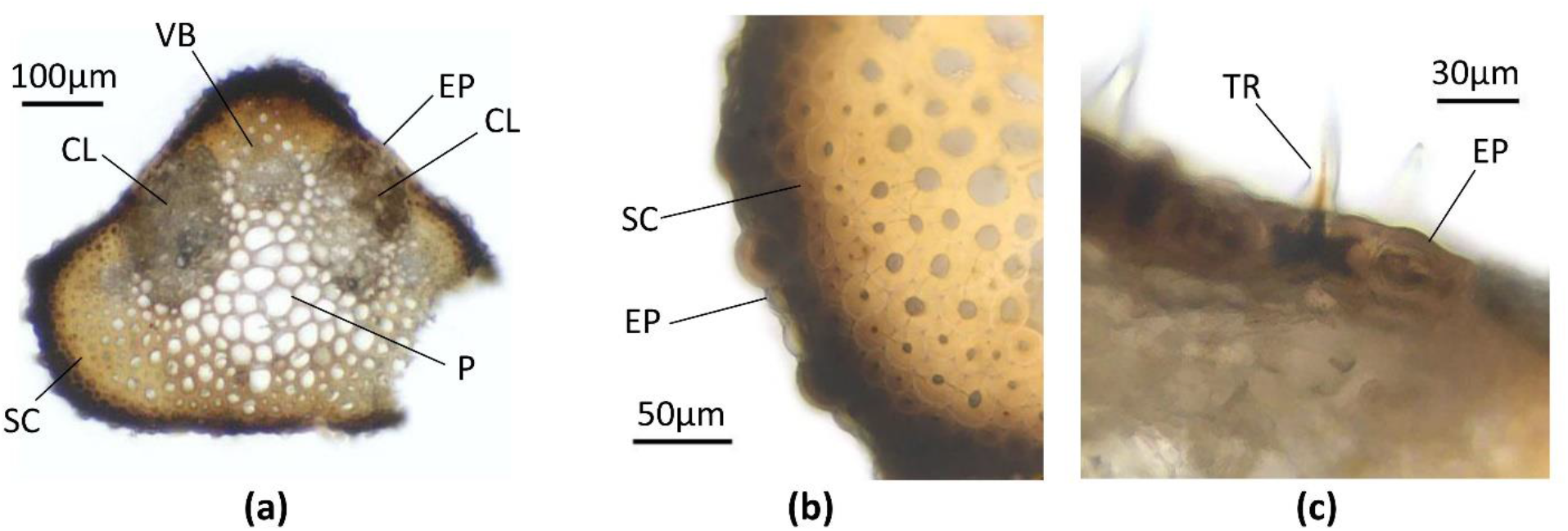
Cross-sections of lemma of dark-colored Persian wheat mature spike, no staining applied. Awn **(a)**, awn at higher magnification **(b)**, lemma broad part **(c)**. VB – vascular bundle, EP – epiderma, CL – chlorenchyma, P – central parenchyma, SC – sclerenchyma, TR - trichome.

## Discussion

Traditionally, dark color of wheat glumes was attributed to anthocyanin content [23]. However, the standard procedure of acidified alcohol extraction suitable for anthocyanins failed to produce colored solution for both dark-spiked wheat husks and sunflower black-colored achenes pericarp. At the same time, alcohol extraction of anthocyanins was successful for tulip perianth, which is a source of pelargonidin 3-rutinoside, cyanidin 3-rutinoside and their acetyl esters in the red tissues, and the cyanidin 3-rutinoside and delphinidin 3-rutinoside in the dark tissues[25], and in blueberry fruit skin, which is a source of various delphinidin, malvidin, petunidin and cyanidin glycosides [27]. Thus, our extraction experiments showed that the dark pigment in the Persian wheat spike is not of anthocyanin origin. Sodium hydroxide extraction, on the contrary, produced colored solutions both in sunflower achenes, which are the proven source of melanin [24], and in Persian wheat dark-colored spike husks. Spectral measurements showed similarity of sunflower and wheat extracts light absorbance. Thus, we can conclude that dark-colored Persian wheat spike husks contain melanin. Histologically, we showed that melanin in wheat spike husks is concentrated in superficial layers of cells – in epidermis and sub-epidermal sclerenchyma, which may imply a protection function of this substance.

On one hand, dark coloration of spikes may be just an evolutionary and agronomically neutral mutation. On the other hand, we can hypothesize its benefits for surviving and reproduction in wild nature for wheat ancestors, and further transmission of this trait to first cultivated landraces. For example, in wild emmer wheat species like *Triticum dicoccoides*, in which spikes at maturity breaks into spikelets and the grain remains within husks, melanin may serve as a protection from insects, making tissues mechanically more rigid or toxic for them. We know that in fungi melanin is essential for long survival of sclerotia and spores in the soil, evidently preventing fungal cell walls from enzymatical degradation by soil microflora [28]. Similarly, the presence of melanin in wild wheat husks could make them more resistant to hydrolytic enzymes of soil bacteria and fungi. Dark coloration may even be a protection from rodents and birds making the spikelets less visible on dark soil surface [1] or acting through color non-preference as birds and rodents do have color preferences [29,30]. There could be other benefits of melanin accumulation for plants. We can assume that melanin, like anthocyanins, may play role in protection of plants from excess of light and damaging ultraviolet radiation [31]. Dark coloration of wheat spikes favors conversion of light into heat. This can possibly speed up the process of maturation [1].

The black color of awns is observed among some durum wheat cultivars [33], while black glumes potentially containing melanin could be found in wheat lines carrying alien genetic material [23,34]. If proved to contain melanin, these wheat genotypes could become an object of practical studies. At the same time, dark coloration of spikes is a rare trait in modern wheat cultivars. That may be a matter of chance as only little genetic diversity has passed from wild species to modern wheat cultivars. Also, it may be just a color preference of the most of farmers and breeders. At last, and most probably, this is a result of selection for higher grain yield, as dark pigmentation of epidermis screens part of light that could be used for photosynthesis. However, melanin containing in spile husks could be beneficial even for modern cultivars, as it may give some resistance against some fungal infections, like *Fusaruim* head blight, as was shown in oats [32]. The potential pests and diseases protection properties of melanin in wheat spikes should be further studied.

## Conclusion

The dark-colored Persian wheat spikes contain melanin, which is concentrated in superficial layers of cells – in epidermis and sub-epidermal sclerenchyma of husks. Protective functions of melanin against wheat pests and diseases should be further studied.

## Footnotes

1 Federal Research Center N. I. Vavilov All-Russian Institute of Plant Genetic Resources

